# Succession of bacteria and archaea within the soil micro-food web

**DOI:** 10.1101/2024.05.08.593117

**Authors:** Mandip Tamang, Johannes Sikorski, Miriam van Bommel, Marc Piecha, Tim Urich, Liliane Ruess, Katharina Huber, Meina Neumann-Schaal, Michael Pester

## Abstract

Bacterivorous nematodes represent numerically abundant bacterial grazers in the soil micro-food web. Their trophic regulation shapes the soil microbiome, but the underlying population dynamics of bacteria and archaea are poorly understood. Here, we followed bulk soil respiration and time-resolved population dynamics (32 days) of bacterial and archaeal species in response to top-down control by a common bacterivorous soil nematode, *Acrobeloides buetschlii*, bottom-up control by resource amendment via maize litter as well as the combination of both. Addition of maize litter significantly increased soil respiration rates, while bacterivorous nematodes shifted the maximum rate of soil respiration from day 12 to day 6. Underlying bacterial and archaeal abundance changes could be separated into five major response types, dominating in different top-down and bottom-up control scenarios. Individual microbial species switched between response types depending on the different scenarios. In-depth analysis of these differential abundance patterns revealed a broad feeding behavior for *A. buetschlii* on dominating populations of gram-negative bacteria (*Acidobacteriota, Bacteroidota, Gemmatimonatoda, Pseudomonadota*) and ammonia-oxidizing archaea (*Nitrososphaerota*), while discriminating against dominant populations of gram-positive bacteria (*Actinobacteriota, Bacillota*). Combined bottom-up control by maize litter and top-down control by nematode grazing caused a succession of soil microbiota, which was driven by population changes first in the *Bacteroidota*, then in the *Pseudomonadota*, and last in the *Acidobacteriota* and *Nitrososphaerota*. This mechanistic understanding of nematode grazing on soil microbiota population dynamics is essential to inform predictive models of the soil food web.

## Introduction

The soil microbiome comprises the most diverse biological communities on Earth, encompassing at least 25% of our planet’s biodiversity [1] with tens of millions of bacterial, archaeal, fungal, and microeukaryotic species [2]. Interestingly, a recent meta-analysis of 237 soils across six continents established that only 2% of bacterial phylotypes (about 500 in total numbers) found in soils consistently accounted for 41% of the soil bacterial communities worldwide. Thus, relatively few bacterial taxa are dominating in numbers across soils globally [3]. The eight most abundant bacterial phyla found in soils (in decreasing relative abundance) are typically represented by heterotrophic members of the *Pseudomonadota* (mainly *Alpha-, Beta- and Gammaproteobacteria*), *Actinobacteriota*, *Acidobacteriota*, *Planctomycetota*, *Chloroflexota*, *Verrucomicrobiota*, *Bacteroidota*, and *Gemmatimonadota* [3, 4]. They are accompanied by the archaeal phylum *Nitrososphaerota* comprising typically 1-5% of the total prokaryotic community [5] and consisting of ammonia-oxidizing chemolithoautotrophs [6, 7].

It is evident that the ecology of the soil microbiome has direct impact on soil ecosystem services, e.g., nutrient cycling, carbon sequestration, and promoting plant growth to name a few [8, 9]. Also, a large share of global soil respiration is caused by the soil microbiome (35 – 69 Pg C/year) [10, 11] because of its inherent role in organic matter decomposition and mineralization [12, 13]. Therefore, the soil microbiome has direct impact on carbon exchange between the land surface and the atmosphere [14, 15]. Three major mechanisms were identified to govern the ecology of the soil microbiome. As a principle regulator, abiotic soil properties such as pH, organic matter content and climate were suggested [16]. In addition, two biotic control mechanisms were identified, particularly at the finer spatial scale. Bottom-up control is exerted by litter input and rhizodeposition, whereas top-down control is driven by predators and viruses [8, 12]. While the importance of bottom-up control, i.e. plant input, is already well recognized and studied [17-22], the role of top-down control by predation has received much less attention [8, 12, 23, 24].

In particular nematodes and protozoa are recognized as regulators of microbial community structure and by driving nutrient mineralization of their microbial prey [25, 26]. Their grazing activity further accounts for the release of ammonium from microbial biomass, estimated as high as 32–38% of the annual N-mineralization in arable land [27]. Thus, eukaryotic grazers link the microbial with the faunal food web in soils [28-30] and hence the energy and matter flow to higher trophic levels. Nematodes constitute the most abundant (5×10^4^ – 27×10^6^ individuals m^-2^) and diverse (up to 200 species m^-2^) multi-cellular organisms in soils [28, 31, 32], with established functional groups at each trophic level, feeding on bacteria, fungi or roots as well as microfauna [33]. Bacterivorous nematodes represent the functionally dominating group within soil nematode communities, typically accounting for >40% of all nematode counts across different terrestrial biomes worldwide [32].

Nematodes feeding on bacteria are often described as generalists that randomly ingest any bacteria by their filter-feeding habit [8]. However, there is increasing evidence that they do differentiate their prey bacteria based on morphology, cell-wall characteristics, mucus production or metabolite concentration [34, 35]. Especially small-sized Gram-negative bacteria that fit the stoma of nematodes were reported to be preferred [34, 36, 37]. Also, a high water content, a high C/N ratio as a proxy for food quality as well as the respiration rate as a proxy for metabolic activity were identified as positive prey selection criteria of bacterivorous nematodes [23, 34, 38]. Differing life-history traits among nematode species are yet another differentiating factor and are commonly expressed as the colonizer-persister (cp) value [39, 40]. Enrichment specialists such as the model nematode *Caenorhabditis elegans* (cp 1) are only active under food-rich conditions as supported by its wide tubular stoma and consumption of food particles through continuous pumping of the oesophagus (pharynx). In contrast, opportunistic nematode species that are common in many soils, such as *Acrobeloides buetschlii* (cp 2), are less sensitive to changes in food type and quantity. In the latter, this goes along with a narrow, muscular stoma and a lower pumping frequency with its pharynx [38-40].

Grazing activities by bacterivorous nematodes alter soil respiration [23, 41, 42] and positively effect organic N mineralization [43-45]. In combination with their feeding preferences, this translates directly into changes of the total abundance and community composition of soil bacteria [41, 46, 47]. However, the underlying population dynamics of individual bacterial and archaeal species are only rudimentarily explored and are typically only described in terms of beta-diversity changes focusing on end-points of the respective experiments. Here, we followed time-resolved population dynamics of bacterial and archaeal species at the absolute abundance level in response to top-down control by a common bacterivorous soil nematode, *A. buetschlii*, bottom-up control by maize litter addition, and the combination of both.

## Materials and Methods

### Experimental design

Farmyard manure-fertilized soil was collected in August 2021. Soil was taken randomly from the upper 20 cm of a long-term field experiment at the Dikopshof research farm (50°48′21″ N, 6°59′9″ E), Germany. Soil and site characteristics are given in the Supplementary Material. After transport to the laboratory, soil samples were air dried at room temperature (RT) for at least 14 days, homogenously mixed, sieved (2 mm mesh), and stored at RT. Prior to experiments, stored soil was pre-incubated for 10 days with 14% (w/v) demineralized water at RT. Thereafter, soil microcosms were set up as described previously [23]. Per microcosm, 50 g dry weight (DW) of pre-incubated soil was maintained at 20°C in the dark under a continuous air-flow with reduced CO_2_ content (3 ppm) and 33% relative humidity. The water content was kept constant at 16%. Further details are given in the Supplementary Material. Treatments of microcosms followed a 2-factorial design: with or without maize litter and with or without *A. buetschlii*, resulting in four treatment combinations (Supplementary Figure 1). Maize litter (IsoLife, Wageningen, The Netherlands) originated from cob, leaf, and stem (milled <2 mm) and was supplied as 0.76 mg (g soil DW) ^−1^, which was roughly equivalent to fourfold of microbial biomass in the used soil. Maize litter was ^13^C-labeled (>97 atom% ^13^C) for carbon flux analyses, which will be described elsewhere (van Bommel et al., in preparation). Nematodes were extracted from monoxenix stock cultures (Supplementary Material) and added at a final amount of 435 ± 35 *A. buetschlii* individuals per microcosm. Each treatment was set up in five replicates per timepoint that were sampled destructively at days 4, 8, 16, and 32. Samples at day 0 were shared in the treatments with and without *A. buetschlii*. Nucleic acid extraction, quantitative PCR as well as 16S rRNA gene amplicon sequencing of total Bacteria and Archaea followed standard procedures detailed in the Supplementary Material.

### Soil respiration

Soil respiration was determined as described previously [23, 48]. In brief: CO_2_ trapped in 1 M KOH-containing vials was determined every 2-4 days. Background CO_2_ was determined by using empty microcosms without soil (blanks). Strontium carbonate (SrCO_3_) was precipitated out of 3 ml CO_2_-containing KOH solution using 3 ml 0.5 M SrCl_2_. The precipitate containing solution was amended with 2 drops of 1% phenolphthalein solution and titrated with 0.1 M HCl until color change. The trapped CO_2_ was calculated according to the following equation:

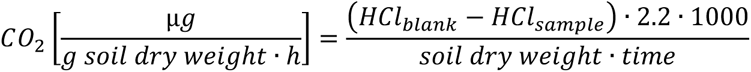

where HCl_blank/sample_ are the volumes of HCl consumed by blanks and samples [ml] and 2.2 is the conversion factor of 0.1 M HCl to 2.2 mg CO_2_. The factor 1000 was used to convert mg into µg. Generalized additive modelling (GAM) of soil respiration rates was done in R package mgcv v. 1.8-41 [49]. GAM model development was performed as recommended previously [50]. Model assumptions were verified as suggested previously [51] by plotting residuals versus fitted values via gratia::appraise() v. 0.8.1.38 [52] (Supplementary Figure 2).

### Alpha and beta diversity of bacterial and archaeal amplicon sequencing variants

Analyses of quality-controlled 16S rRNA gene amplicon sequencing variants (ASVs) and qPCR data were performed in R version 4.2.2 [53]. Graphs were generated using R package ggplot2 v.3.4.4 [54]. Sampling coverage [55] was estimated on the basis of the Qiime2-processed ASV table in R package iNEXT v. 3.0.0. [56]. Coverage equaled for all samples 1 indicating close to complete sampling coverage. As a consequence, no downstream normalization based on rarefaction was applied. If indicated, relative abundances were transformed to absolute abundances by multiplying with the respective total 16S rRNA gene copy numbers as obtained by qPCR. Alpha diversity indices based on Hill number analysis were obtained on absolute abundance transformed ASV tables using the R package hilldiv v.1.5.1 [57]. The alpha-gambin value as a parameter describing species abundance distributions [58] was obtained on read count ASV tables using the R package gambin v. 2.5.0 [59]. A higher alpha-gambin value reflects a flatter species abundance distribution, indicating a more even distribution of species abundances. To test for significant alpha diversity changes an analysis of variance (ANOVA) was performed using the R package stats v. 4.2.2. Significant ANOVA results were subjected to post-hoc comparisons of group means using the R packages multcomp v.1.4-25 [60] and sandwich v.3.0-2 [61, 62] as described previously [63]. Distance matrices for beta-diversity analysis were obtained using the R package phyloseq v. 1.42.0 [64]. Nonmetric Multidimensional Scaling (NMDS) ordination and variance partitioning from distance matrices were done using the R package vegan v. 2.6-4 [65].

### Classification of ASV abundance changes into response types

Classification of time-resolved ASV abundance changes into response types was based on numerically dominant ASVs as defined by Hill number ^2^*D* [66]. Briefly: Hill number ^2^*D* is an established measure in ecology on the effective number of dominant taxa in a sample and is derived from the inverse of the Simpson index [66]. Its big advantage is that it considers both, species richness and evenness. As both may differ between different samples, the Hill number concept should not be confused with the simpler cutoff of 0.1% relative abundance previously introduced to microbial ecology to differentiate low-abundance species (rare biosphere) from so called “abundant” species [67-69]. In the Hill number concept, species richness (^0^*D*), abundant species (^1^*D*), and dominant species (^2^*D*) are differentiated and their respective cutoff is depending on the species richness and evenness in a given sample.

For response type analysis, ASVs had to be dominant in at least one of the 54 samples. In addition, dominant ASVs had to be detected in at least two out of three replicates per time across all four experimental treatment combinations. This resulted in a subset of 359 dominant ASVs, which had a relative abundance of 0.0035 to 7.9% (mean = 0.19%; median = 0.09%) across all samples. To discern response ASVs among this subset, the latter were subsampled based on their absolute abundance range, i.e. minimum to maximum at any given timepoint. Following the Pareto principle, only ASVs were considered for downstream analysis which summed absolute abundance ranges across all treatments represented 80% of the total absolute summed abundance ranges of all subset ASVs. This resulted in 221 dominant response ASVs, which were further grouped into response types. To foster comparability across all response ASVs, absolute abundances of each ASV were first scaled to a mean = 0 and a standard deviation = 1 using the R package stats v. 4.2.2. Thereafter, scaled response patterns were subjected to k-means clustering. An initial pre-screening analysis was performed using three different clustering methods as implemented in the R package factoextra v. 1.0.7 (http://www.sthda.com/english/rpkgs/factoextra). The method “gap_stat” was used with the additional arguments nstart = 25 and nboot = 50. Both, the “wss” (within cluster sums of squares) and the “silhouette” method suggested four optimal clusters, whereas the “gap_stat” (gap statistic) suggested nine optimal clusters. As a result, the analysis of response types was initiated with a k-means clustering using nine clusters. Visual inspection identified highly similar response types, which were further grouped into a final set of five different response types, which was justified by the suggested 4-9 optimal clusters.

### Data availability

All amplicon sequences were deposited at the Sequence Read Archive at National Center for Biotechnology Information (NCBI) under the bioproject number PRJNA1092885.

## Results

### Bacterivorous nematodes speed up soil respiration

Soil microcosms were set up to examine biotic controls of the bacterial and archaeal community in a typical agricultural soil. Top-down control by grazing was investigated by addition of the bacterivorous nematode *A. buetschlii*. Bottom-up control by plant input was investigated by addition of maize litter (Supplementary Figure 1). Total nematode densities were regularly checked for all treatment combinations. In the treatments with *A. buetschlii* addition, total nematode densities were initially 5.0 ± 1.5 individuals (g soil DW) ^−1^, which corresponded well to the theoretical value of 8.7 added *A. buetschlii* ind. (g soil DW) ^−1^. At day 8, nematode counts increased slightly to 8.6 ± 4.5 and 7.8 ± 2.1 ind. (g soil DW) ^−1^ with and without added maize litter, respectively. At day 32, nematode counts increased substantially to a total of 20.5 ± 7.0 and 14.1 ± 6.7 ind. (g soil DW) ^−1^ with and without added maize litter, respectively. In the treatments without *A. buetschlii* addition and no maize amendment autochthonous nematode counts were much lower, ranging between 0.3 ± 0.0 and 2.2 ± 2.1 to ind. (g soil DW) ^−1^ across analyzed dates. With maize but without *A. buetschlii* inoculation, autochtonous nematode counts were still low with 1.2 ± 0.3 (g soil DW) ^−1^ at day 8 but increased until day 32 to 16.4 ± 3.5 ind. (g soil DW) ^−1^. Addition of *A. buetschlii* clearly shifted the maximum rate of soil respiration to an earlier time point (from day 12 to day 6, Figure 1) and homogenized the oscillation of CO_2_ release from soil. Presence or absence of added nematodes also explained best dynamics of soil respiration over the examined 32 days (Supplementary Table 1). In contrast, addition of maize litter significantly (*p* < 0.05) increased soil respiration rates, whereas the addition of nematodes or the combined effect of both had no such effect (Supplementary Table 2).

**Figure 1.**
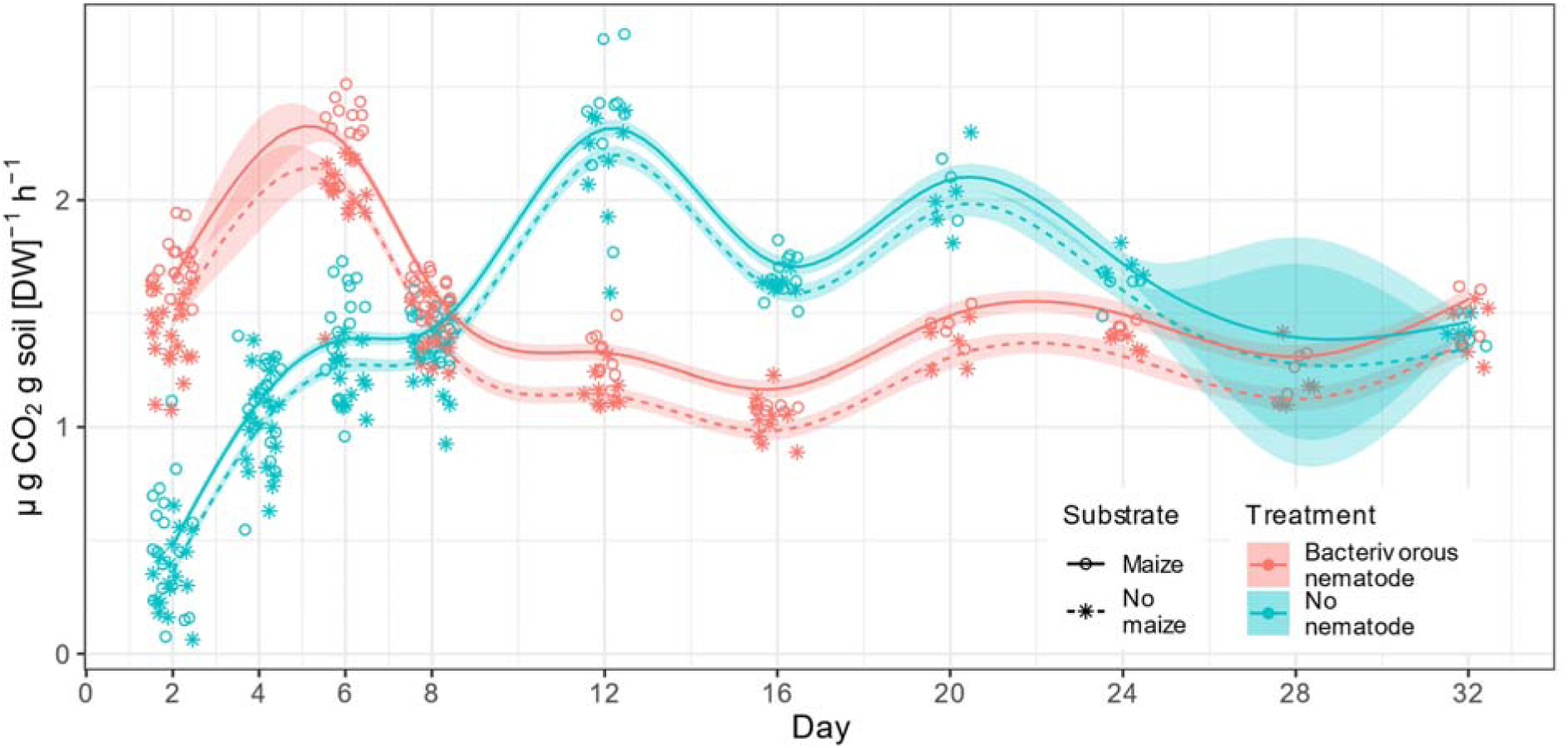
Soil respiration rate in response to addition of the bacterivorous nematode *A. buetschlii* and maize litter. Individual measurements are given as symbols under the given treatments. In addition, a generalized additive model (GAM) was fitted to the measured values and shown as continues solid or dashed lines. Details of the GAM are given in Supplementary Tables 1 and 2 as well as in Supplementary Figure 1. The standard error is shown as shades next to the GAM.

### Grazing and substrate availability alter bacterial and archaeal community composition

Total bacterial and archaeal abundances, ASV richness, and the number of dominant ASV as defined by Hill number ^2^*D* [66] were on average 2.35 ± 0.53 × 10^8^ 16S rRNA gene copies per gram dry soil, 2.06 ± 0.39 × 10^3^ ASVs, and 1.54 ± 0.38 × 10^2^ ASVs at the onset of the experiment, respectively (Supplementary Figure 3). Total bacterial and archaeal abundances changed only significantly over time when both the bacterivorous nematode and maize litter were added (Supplementary Tables 3, 4). Species richness, the number of dominant ASVs and the rank abundance distribution of the bacterial and archaeal community did not change significantly throughout the incubation in any of the incubation types (Supplementary Tables 3, 4).

While alpha diversity metrics stayed rather stable throughout the experiment, there was a pronounced effect of both the bacterivorous nematode and maize litter addition on the overall community composition of bacteria and archaea (Supplementary Figure 4). Most of these community changes occurred within the first 16 days of the experiment, while there were little changes in the remaining 16 days (Supplementary Figure 5). Variance partitioning of Bray-Curtis distances could explain 35% of the variance in the data with nematode treatment having the largest effect, followed by incubation time and maize litter addition (Figure 2). When considering also the phylogenetic relatedness of ASVs as in weighted unifrac, 40% of the variance in the data could be explained. Here maize litter addition had by far the largest effect, followed by nematode treatment and incubation time (Figure 2). In combination, this showed that phylogenetically related ASVs were stronger differentially affected by nematode treatment, while the addition of maize litter stronger influenced distantly related ASVs.

**Figure 2.**
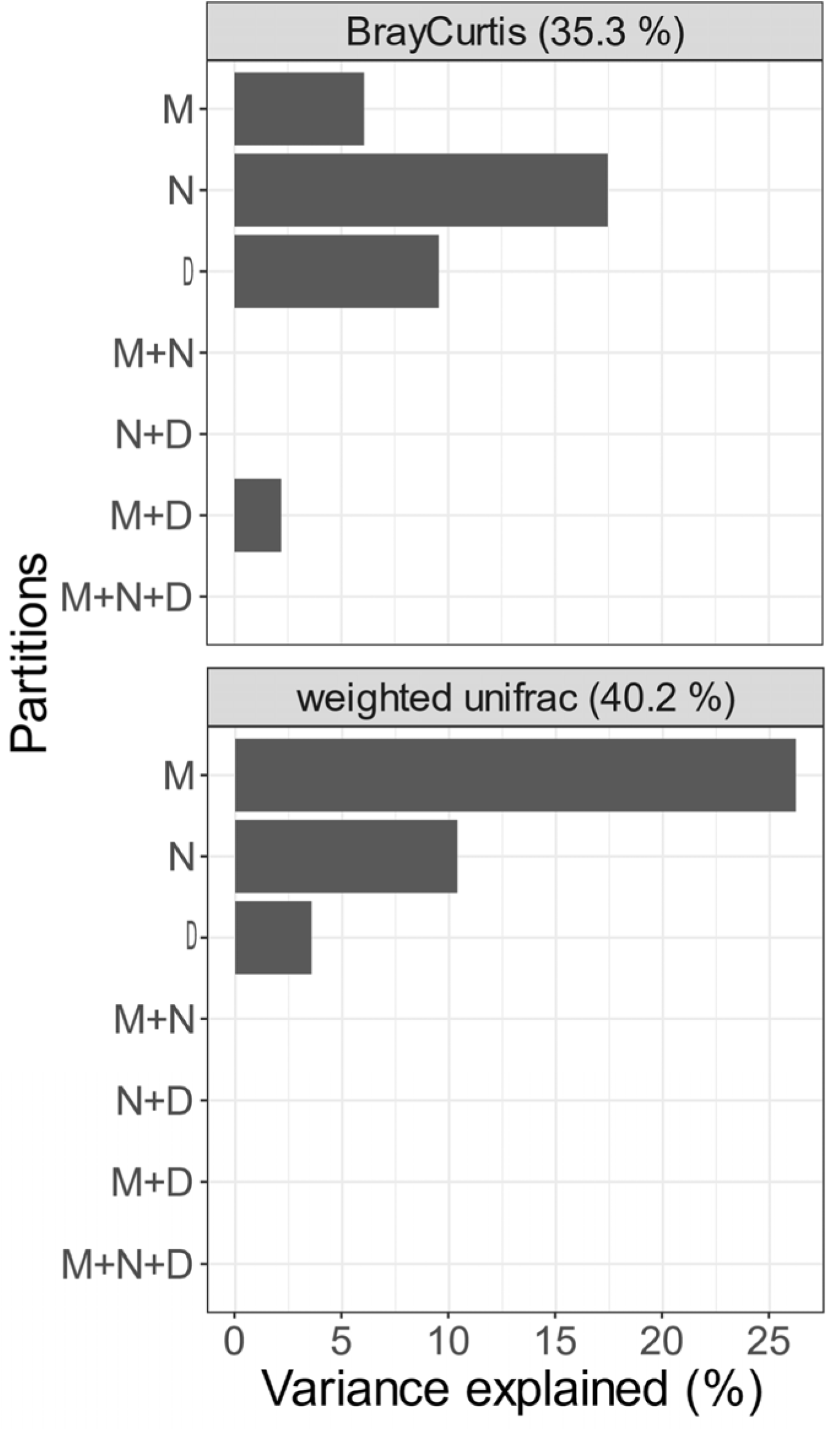
Variance partitioning of beta diversity changes of the bacterial and archaeal community. M – Maize litter addition, N – Nematode treatment, D – incubation time. Interactions of individual explanatory variables are indicated with the plus sign.

### Five ways bacteria and archaea numerically respond to top down and bottom up control

Abundance changes of individual ASVs in response to nematode treatment, maize litter addition, and the combination of both were analyzed by grouping ASVs into time-resolved response types. The focus was put on dominant ASVs as defined by Hill number ^2^*D* (see Material and Methods for a detailed description) as these are considered to drive the major ecosystem functions. This approach was preferred over classical differential abundance testing since it considered (i) temporal dynamics across multiple time points (ii) focused on populations with major absolute (and not relative) abundance changes, and (iii) did not suffer from *p*-value inflation due to massive parallel, pairwise testing and the resulting sensitivity loss of detecting biologically meaningful population changes.

Five different response types at the population level were identified, which were labeled from A to E (Figure 3A). In the presence of the bacterivorous nematode (N) and under maize (M) litter addition (+N/+M), response type E was dominating in terms of number of ASVs and contribution to overall abundance changes of total bacteria and archaea (Figure 3 B,C). It was characterized by an intermediate abundance peak at day 8 and decline to starting point values towards day 32. In the parallel treatment without nematode but maize litter present (−N/+M), response type D became dominant in terms of ASV numbers and contribution to overall abundance changes. Interestingly, response type D was very similar to response type E but the intermediate abundance peak was shifted forward to day 4 (Figure 3 B,C). In the treatment with nematode but without maize litter (+N/−M) yet two other response types, type A and B, were dominating. Response type A was characterized by a steady decline in abundance over time, while response type B was characterized by an intermediate abundance minimum at day 8 and a recovery to starting point values at day 32 (Figure 3 B,C). In the control treatment without nematode and maize litter addition (−N/−M), response type D was dominating, as was the case in the treatment without nematode but with maize litter (−N/+M). In addition, response type C, which is characterized by a steady abundance increase over time, became dominant towards the end of the incubation as well (Figure 3 B,C).

**Figure 3.**
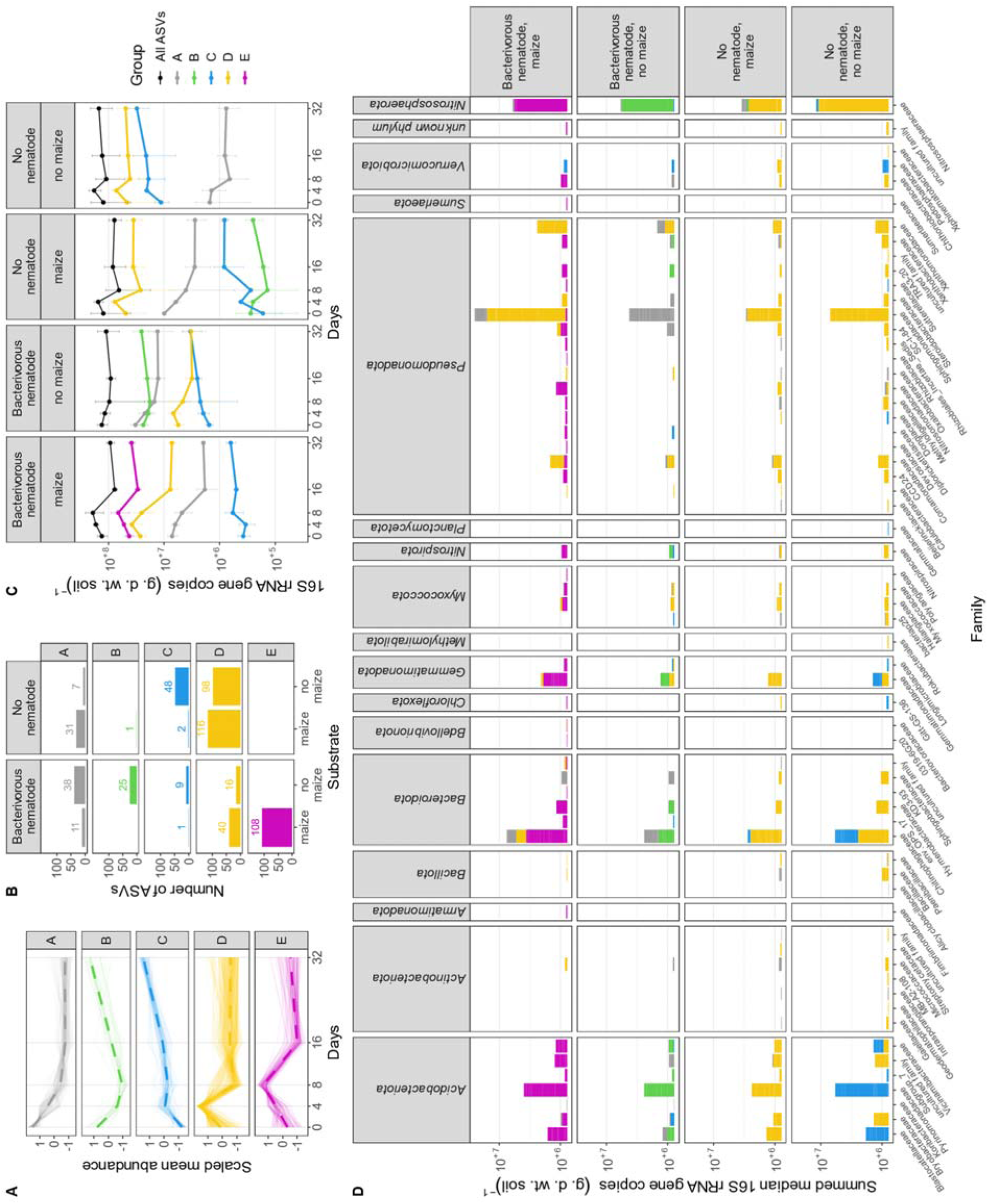
Classification of ASV abundance changes into response types across the different treatment combinations of the bacterivorous nematode *A. buetschlii* (top-down control) and maize litter addition (bottom-up control). (A) Response types as identified by k-means clustering of absolute abundances scaled to a mean = 0 and a standard deviation = 1. For each response type, thin solid lines indicate scaled abundance changes of representing ASVs and the dashed thick line indicate the respective median. (B) Number of dominant response ASVs belonging to the individual response types under the different treatment combinations. (C) Contribution of response types to overall absolute abundance changes of the total bacterial and archaeal community. The mean ± standard deviation is provided for 2-3 replicates. (D) Summed median abundances of ASVs across all incubation timepoints. ASVs were grouped at the family level and according to the different response types under the different treatment combinations.

### Soil bacteria and archaea switch between response type depending on treatment

The +N/+M treatment was clearly dominated by response type E (abundance maximum at day 8). This response type was spanning ASVs across 13 phyla with representatives of the *Acidobacteriota* (families *Blastocatellaceae* and *Pyrimonadaceae*), *Bacteroidota* (*Chitinophagaceae*), *Gemmatimonadota* (*Gemmatimonadaceae*), and *Nitrosphaerota* (*Nitrososphaeraceae*) dominating (Figure 3D). With the exception of the archaeal *Nitrososphaerota* (see below), most other dominant ASVs represented bacterial groups with a typically heterotrophic lifestyle. Interestingly, ASVs representing response type E under the +N/+M treatment switched to different response types under other treatment combinations. For example, *Acidobacteriota* ASVs with large average abundances switched to an earlier abundance peak at day 4 in the −N/+M treatment (response type D), to an abundance decline towards day 8 and subsequent recovery (response type B) in the +N/−M treatment, and to a steady abundance increase (response type C) in the −N/−M treatment (Figure 4, Supplementary Table 5). The second most important response type in the +N/+M treatment was response type D. In contrast to response type E, this response type was clearly dominated by *Pseudomonadota* ASVs, in particular representatives of the *Sphingomonadaceae*, *Xanthomonadaceae*, and *Comamonadaceae* (Figure 3D). Here, the response pattern stayed quite stable throughout the different treatment types with the only exception being the +N/−M treatment, where a constant decline (response type A) was observed for some of these dominant ASVs (Figure 4, Supplementary Table 5). In total, 56 of such response type patterns could be observed in our data set (Supplementary Figure 6) with 13 of those representing >80% of the summed median abundances of all dominant ASVs that were grouped into response types (Figure 4). The four most prominent response type patterns in terms of summed abundance had in common that their representing ASVs showed a decrease in abundance in the first eight days in the +N/−M treatment (response type A and B) but showed the opposite behavior with an abundance peak at day 4 in the −N/+M treatment (response type D) (Figure 4). They mainly differentiated in their response to the +N/+M treatment and to a minor extent in their behavior in the −N/−M control treatment (Figure 4).

**Figure 4.**
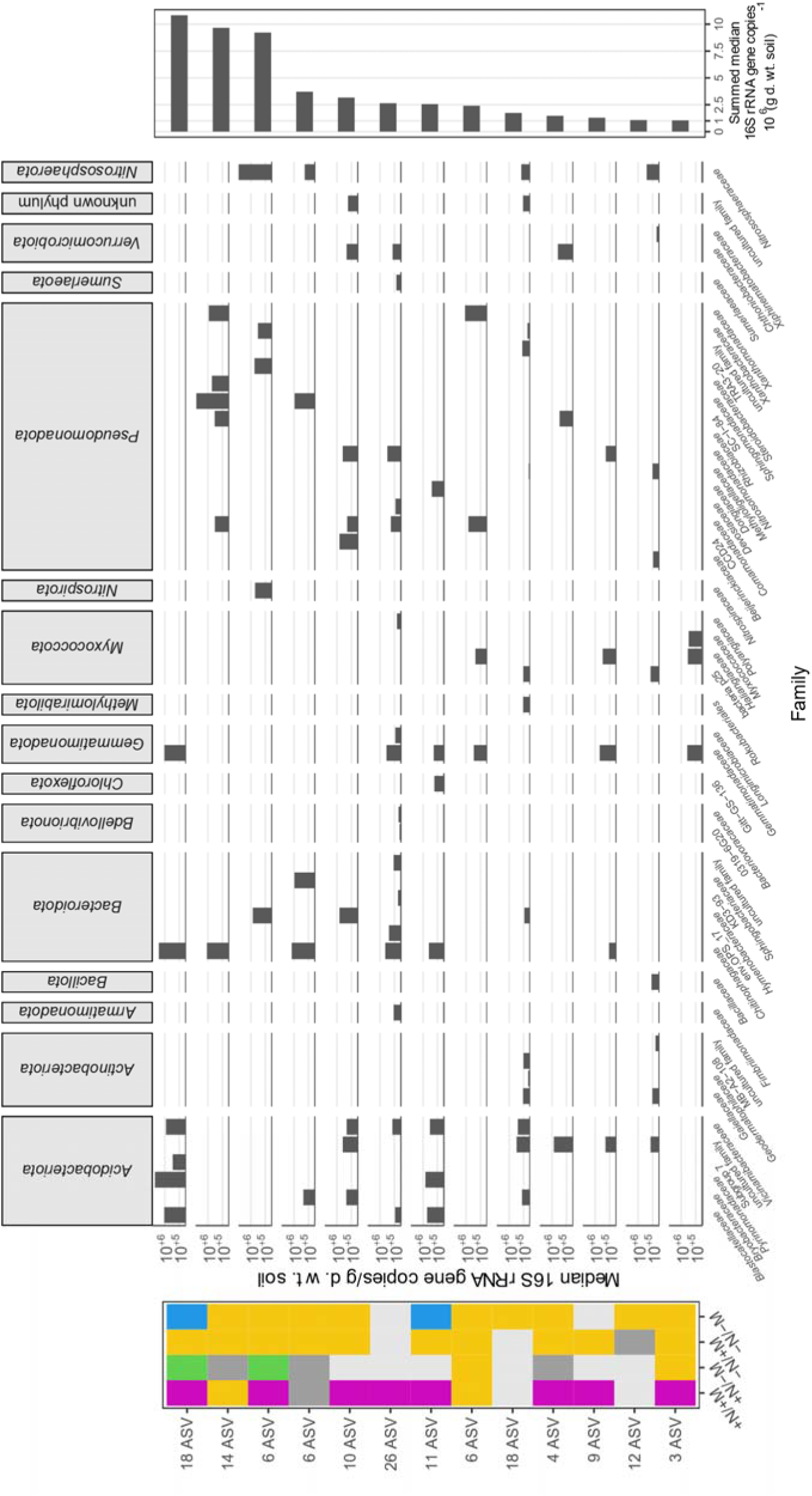
Response type changes of dominant response ASVs across all four treatments. The color code corresponds to the response type shown in Fig. 3. The number of ASVs belonging to a certain response type pattern is indicated to the left. The taxonomic affiliation of respective ASVs and their median absolute abundance across all treatments and time points is shown in the center. The median total abundance correlated highly significantly with the mean total abundance (Pearson’s *R^2^*= 0.997, *p*-value < 2.2 × 10^-16^). The median total abundance was chosen to indicate the base line of the total abundance of the respective ASV throughout the entire experiment. The summed median abundance of all ASVs belonging to the same response type pattern is shown to the right. Only response type patterns integrating over the majority of dominant response ASVs, i.e. 80.9 % of all summed median abundances, are shown. Summed median abundances of all response type patterns are shown in Supplementary Figure 5 and the respective ASVs are given in Supplementary Table5. M – maize litter , N – nematode, + and – signs indicate presence and absence, respectively.

An outstanding group of microorganisms with a specialized lifestyle were bacteria and archaea belonging to the functional guild of nitrifiers, i.e. chemolithoautotrophic microorganisms converting ammonia via nitrite to nitrate. Especially *Nitrososphaerota* ASVs, which all belonged to typical soil ammonia oxidizing archaea within the *Nitrososphaeraceae* [6, 70, 71] constituted one of the most abundant and responsive groups. While largely belonging to the dominant response type E in the +N/+M treatment, they clearly switched to response type B when maize litter was lacking. If the nematode was lacking, they switched to response type D irrespective of maize litter addition (Figure 3D, Figure 4). The same response pattern dynamics were observed for *Nitrospirota* ASVs affiliated to the genus *Nitrospira* that comprises typical soil nitrite-oxidizing bacteria and complete ammonia oxidizers [72] (Figure 3D). Ammonia oxidizing bacteria affiliated to the *Nitrosomonadaceae* (*Pseudomonadota*) constituted the third group within this functional guild, yet at smaller population size as compared to the *Nitrososphaeraceae* and *Nitrospira* ASVs. As the other nitrifiers, they belonged to response type E in the +N/+M treatment and most shifted to response type D if just the nematode was omitted (−N/+M). In contrast to the other nitrifiers, the majority of *Nitrosomonadaceae* ASVs did not show relevant abundance changes in the +N/−M and −N/−M treatments (Figure 4, Supplementary Table 5).

## Discussion

### The broad food range of a bacterivorous nematode spans two domains of life

Adult nematodes of *A. buetschlii* and other bacterivorous species can consume about 10^5^−10^6^ bacterial cells daily [73]. Their prey is essentially sucked in by the pumping action of the pharynx and hence their feeding behavior can be regarded as filter-feeding [74, 75]. Therefore, the primary selecting factor of their food choice is constrained by the width of the stoma that determines the maximum size of prey bacteria ingested [75]. Any bacteria larger in size or growing in dense micro-colonies or biofilms will thus efficiently reduce their palatability [75]. In addition, binary interaction assays established that bacterivorous nematodes can display food preferences among bacteria of the preferred size class [23, 34, 36-38] and thus modulate the composition of bacterial soil communities by their grazing activity [41, 46, 47]. These pioneering insights were restricted to bacteria easy to grow in the laboratory. However, major bacterial (e.g., *Acidobacteriota*, *Gemmatimonadota*) and archaeal (*Nitrososphaerota*) taxa dominating in soil were so far neglected.

Our experimental setup allowed us to decipher population dynamics of dominant bacterial and archaeal species-level ASVs exposed to grazing by a typical bacterivorous soil nematode, *A. buetschlii.* The combination of nematode population development, soil respiration dynamics in the presence of *A. buetschlii* (Figure 1), the time-resolved magnitude of bacterial and archaeal community changes (Supplementary Figure 5), and the population response type analysis of bacterial and archaeal ASVs (Figure 3) showed that the first eight days of the experiment were decisive to understand predator-prey effects. Here, comparison of the isolated effect of the added nematode (+N/−M treatment) to treatments without nematode addition (−N/+M; −N/−M) allowed us to identify species-level taxa that clearly declined in population size (response types A and B) in the presence of the nematode but benefited from the absence of the nematode (response types C and D) irrespective of maize litter addition. It was evident that a selection of the most abundant bacterial ASVs in the analyzed soil were negatively affected by nematode addition. Specifically, they were represented in large parts by the four most prominent response type patterns observed in terms of summed median abundance (Figure 4). To a minor extent, dominant ASVs grouped in less prominent response type patterns contributed as well (Supplementary Figure 6, Supplementary Table 5). Although we cannot differentiate between direct and indirect effects of grazing, it is likely that *A. buetschlii* was feeding on major populations of very abundant gram-negative bacteria including the very prominent groups of *Acidobacteriota*, *Bacteriodota*, *Gemmatimonadota*, and *Pseudomonadota*. At the same time, *A. buetschlii* clearly discriminated against abundant populations of gram-positive *Actinobacteriota* and *Bacillota* and selected gram-negative ASVs, e.g., within the *Comamonadaceae* and *Xanthomonadaceae* (Figure 4). This confirms that bacterivorous nematodes do in parts display feeding preferences also in their natural environment as already indicated by targeted experiments with model bacteria [34, 36, 37].

The very abundant group of ammonia-oxidizing archaea (*Nitrososphaeraceae* ASVs) and their syntrophic partners, the nitrite-oxidizing bacteria (*Nitrospira* ASVs), were negatively affected in population size as well when comparing the isolated effect of the nematode (+N/−M treatment) to treatments without nematode addition (−N/+M; −N/−M). Bacterivorous nematodes are well known to enhance soil N mineralization rates and thus provide excess ammonia as the primary substrate for the nitrifiers mentioned above [43, 44, 76-78]. As a consequence, one should expect population increases of these nitrifiers or neutral effects [25]. The fact that these prominent nitrifier groups declined in abundance over the first eight days and only recovered afterwards (response type B) in the +N/−M treatment indicates that they served as primary food source of *A. buetschlii* as well. This palatability towards filter-feeding would agree well with the generally small coccoid cell size of species belonging to the abundant archaeal *Nitrososphaeraceae* [≤ 1.1 µm, 79, 80] and the slightly larger and spiral-shaped bacterial *Nitrospira* [0.2–0.4 × 0.9–2.2 µm, 81].

### Top-down control induced succession of bacteria and archaea under substrate availability

The simultaneous treatment with the bacterivorous nematode and maize litter (+N/+M treatment) added another layer of complexity to our experimental system. It now involved both a top-down and a bottom-up control at the same time, which is a scenario close to natural settings in agricultural soils after harvest. ASVs that were susceptible to nematode grazing alone (as described above) split now into three successional groups:

1. ASVs in succession group I declined in abundance in both nematode treatments (response type A), with and without added maize litter, and thus did not benefit from the applied bottom-up control. It was composed of 9 ASVs, which constituted in sum 4.5 % of the overall bacterial and archaeal community at the onset of the experiment (Figure 5A). Within the +N/+M treatment, they were mainly dominated by representatives of the *Bacteroidota* as deduced from their mean absolute abundance across all time points (Figure 5B).
2. ASVs in succession group II shifted from population declines under the +N/−M treatment (response types A and B) to an intermediate abundance peak at day 4 (response type D) in the +N/+M treatment. This indicates that these ASVs benefited from substrate addition (bottom-up control) despite simultaneous grazing (top-down control) until day 4, which marked the turning point of a stronger top-down control. Succession group II was composed of 16 ASVs, which constituted 13.8 % of the overall bacterial and archaeal community at the onset of the experiment (Figure 5A). They were heavily dominated by *Pseudomonadota* in terms of mean absolute abundance throughout the +N/+M treatment (Figure 5B).
3. Interestingly, there was a third succession group, where the turning point from stronger bottom-up to stronger top-down control was extended to day 8. Succession group III was composed of even more ASVs (34 in total), which constituted 18.6 % to the overall bacterial and archaeal community at the onset of the experiment (Figure 5A). These 34 ASVs were spanning 7 different phyla with representatives of the *Acidobacteriota* (domain *Bacteria*) and *Nitrososphaerota* (domain *Archaea*) dominating in terms of mean absolute abundance throughout the +N/+M treatment (Figure 5B). In summary, this revealed that nematode grazing under simultaneous substrate availability induces a succession of major bacterial and archaeal soil taxa within the soil micro food-web.

**Figure 5.**
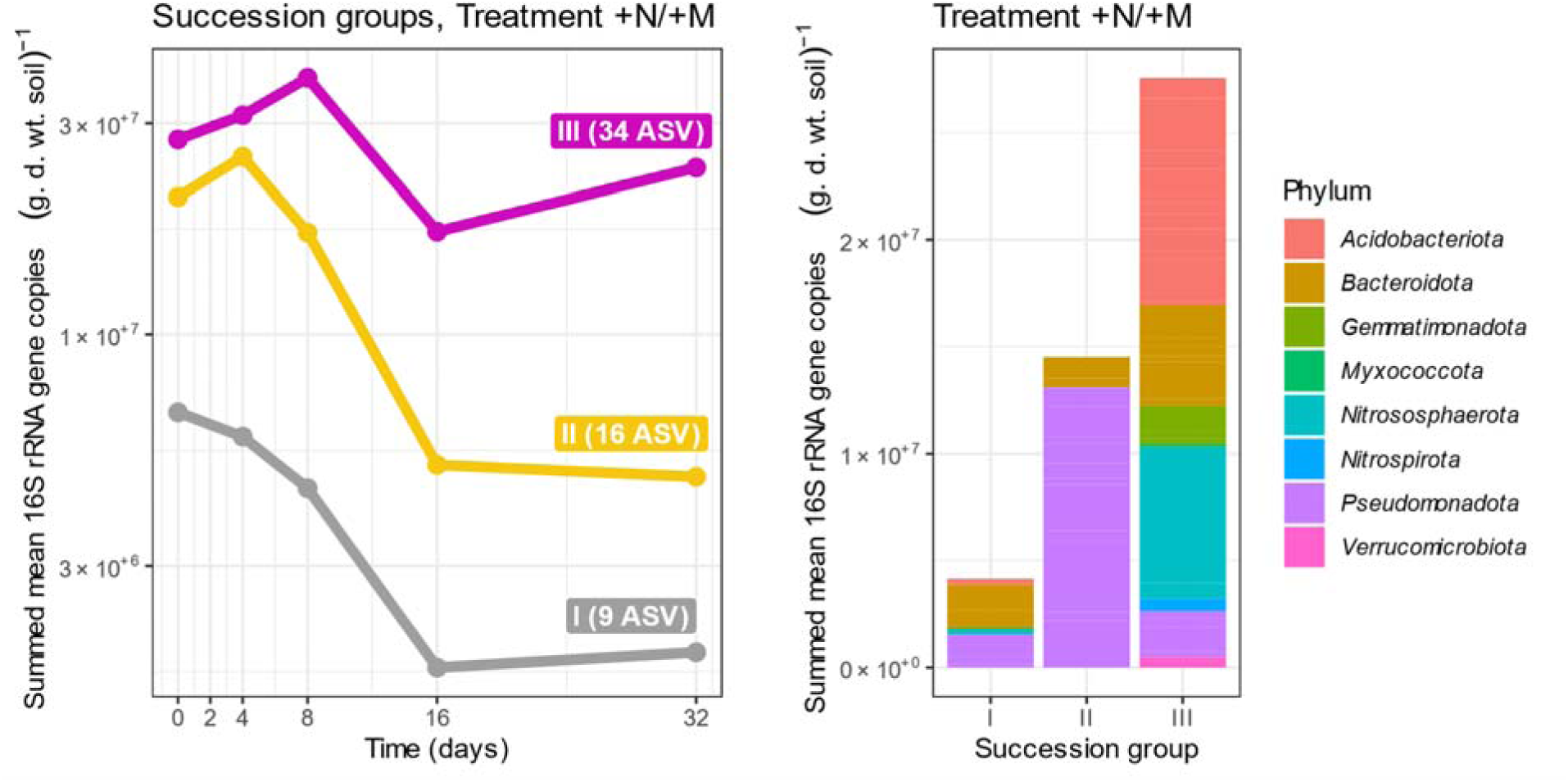
Succession of dominant bacterial and archaeal response ASVs under top-down control by nematode grazing and simultaneous bottom-up control by maize litter addition. Succession groups represent ASVs that were susceptible to nematode grazing (abundance decline within the first eight days in the +N/−M but not in the −N/+M and −N/−M treatments) but split into response types A (succession group I), response type D (succession group II) and response type E (succession group III) in the +N/+M treatment according to Fig. 3. Color codes correspond to Figure 3. (A) Temporal dynamics of succession groups. (B) Affiliation and mean 16S rRNA gene copy numbers of corresponding ASVs across all time points of the +N/+M treatment. The mean was used to indicate the baseline abundance across all time points.

## Conclusion

Soil respiration constitutes a major C flux between the land surface and the atmosphere [14, 15]. Nematode grazing clearly speeded up soil respiration and as a consequence the underlying C flow within soil. This was centered in a broad feeding behavior on dominant populations of gram-negative bacteria and ammonia-oxidizing archaea, while discriminating against dominant populations of gram-positive bacteria. Under simultaneous bottom-up control by maize litter addition, top-down control by nematode grazing caused a succession of microbial groups. This succession was likely driven by a combination of different microbial resource use efficiencies, specialization for substrates with different degradability, and grazing-induced release of degradation products such as ammonia. The latter emphasizes the importance of the micro-food web for nitrogen-use efficiency in soils, impacting ammonia-to-nitrate conversion by nitrifier activity with subsequent nitrate leakage to groundwater and feedbacks on plant productivity [82]. Our temporally resolved succession model provides a first understanding of nematode grazing down to the bacterial and archaeal species level. It lays conceptual grounds for model approaches resulting in predictive ecology of soils benefiting a sustainable agriculture.

## Acknowledgements

We are grateful for the technical support by Morten Streblow, Gesa Martens and Petra Büsing.

## Contributions

The study was conceived and designed by MiPe, MNS, KH, TU, and LR. Experiments and data analysis were conducted by MT, MvB, MaPi, and JS. MiPe and MT wrote the manuscript. All authors edited and approved the manuscript.

