## Supplementary Material for "Succession of bacteria and archaea within the soil micro-food web"


#### Supplementary Methods

##### Soil

Farmyard manure-fertilized soil was collected from the long-term fertilizer field experiment at the Dikopshof research farm (50°48’21” N, 6°59’9” E), which is maintained by the University of Bonn, Germany. The site is located at 62 m above sea level with an average annual temperature and precipitation of 10.5°C and 688 mm, respectively. The dominant soil type is Haplic Luvisol composed of 68.9% silt, 15.9% sand, and 15.1% clay. The soil has a pH (0.01 M $CaCl_2$) of 6.3, a total organic carbon content of 0.74%, and a total nitrogen content of 0.08%. The Dikopshof agricultural site is managed as a 5-year crop rotation of sugar beet (*Beta vulgaris* L.), winter wheat (*Triticum aestivum* L.), winter rye (*Secale cereale* L.), Persian clover (*Trifolium resupinatum* L.), and potatoes (*Solanum tuberosum* L.). On the plots sampled, farmyard manure has been applied annually as sole fertilizer at a rate of 20 tons per ha and crop [1].

##### Nematode

Monoxenix stock cultures of the bacterivorous nematode *A. buetschlii* were maintained at 15°C on potato dextrose agar (Carl Roth, Karlsruhe, Germany) inoculated with the ascomycete *Chaetomium globosum*.

The agar and the compounds secreted by *C. globosum* served as food for *A. buetschlii*. For grazing experiments, *A. buetschlii* individuals were extracted from plates at 20°C for 24 h using a modified wet funnel method after Baermann [2]. Extracted nematodes were surface sterilized with 1 mL 0.01% $HgCl_2$-solution (w/v) for 3 min, followed by washing with 3 mL autoclaved mineral water (Volvic®, Danone waters GmbH, Frankfurt, Germany) for three times. Thereafter, nematodes were stored overnight in autoclaved mineral water at 4°C before the onset of the experiment. The nematode density was determined by light microscopy using 40-fold magnification (Olympus CHT, Olympus optical, Tokyo, Japan). Finally, 1 ml solution consisting of 435 ± 35 *A. buetschlii* individuals was added to respective microcosms.

#### Microcosm setup

The experimental design followed a setup described in detail in Richter et al [3]. In brief: per microcosm, 50 g dry weight (DW) of pre-incubated soil was placed in a 100 ml crimp neck flask (Rotilabo®, Carl Roth) and sealed air tight with crimp caps (Rotilabo®, Carl Roth). Each microcosm was provided continuously with ambient air at a rate of approximately 275 ml min$^{-1}$ using a vacuum pump (Sera® air 550 R plus, Heinsberg, Germany). The inflowing air was first humidified by passing through distilled water and filter-sterilized by a 0.2 µM filter (Sartorius minisart®, Göttingen, Germany) before entering the microcosm. The outflowing air was passed through a separate vial containing 15 ml of 1 M KOH to trap $CO_2$ produced in the microcosms. Microcosms were maintained in a growth chamber (PERCIVAL® E−41L2, Percival Scientific, Perry, Iowa, USA) at 3 ppm $CO_2$, 33 % relative humidity and 20°C in the dark. The water content within microcosms was kept at 16% by regularly injecting autoclaved mineral water with a disposable sterile syringe (Omnifix® 10 ml, Braun, Melsungen, Germany).

#### Nucleic acid extraction and quantitative PCR

Microcosms were sampled destructively for their soil content at days 0, 4, 8, 16 and 32. Soil samples were frozen immediately in liquid nitrogen ($N_2$) and stored at −80°C. Three out of five soil sample replicates per time point and substrate/nematode treatment combination were randomly selected and used in downstream analyses, resulting in a total of 54 samples. DNA was extracted from approx. 2 g of soil using the RNeasy PowerSoil Total RNA kit in combination with the RNeasy PowerSoil DNA elution kit (Qiagen, Hilden, The Netherlands). DNA concentrations were determined fluorometrically using a Qubit 4 (Thermo fisher scientific, Waltham, Massachusetts, USA).

Quantification of total bacterial and archaeal 16S rRNA genes was done as described previously [4] using a quantitative PCR (qPCR) assay. The assay was based on the universal primer pair 1389F (5′-TGYACACACCGCCCGT-3′) and 1492R (5′-GGYTACCTTGTTACGACTT-3′) and conducted in 20 µl reactions on a qTOWER$^3$G (Analytik-Jena, Jena, Germany). PCR reactions consisted of 10 µl of lightcycler ® 480 SYBR Green I master mix (Roche, Penzberg, Germany), 0.4 µl of each forward and reverse primers (10 µM), 5 µl of template DNA (1.6 – 6.5 ng), and 4.2 µl of PCR-grade DNA-free water (Molzym GmbH and Co. KG, Bremen, Germany). After testing a dilution series of template DNA, 1:50 dilutions were used to avoid partial PCR inhibition by co-extracted compounds. An amplified 16S rRNA gene fragment of *Escherichia coli* strain K12 cloned into the pCR 4-TOPO vector (Invitrogen, Waltham, Massachusetts, USA) was used as standard curve in the range $10^2$ to $10^7$ copies µl$^{-1}$ ($R^2$ = 0.99). Amplification was carried out using an initial denaturation at 95°C for 10 min followed by 46 cycles of denaturation at 95°C for 30 sec, annealing at 52°C for 30 sec and elongation at 72°C for 30 sec. PCR efficiencies were on average 90.0 ± 2.8. Specificity of qPCRs was checked by running a melting curve after each qPCR run.

#### 16S rRNA gene amplicon sequencing and sequence processing

Total bacterial and archaeal 16S rRNA genes were amplified using a procedure described before [5] with minor modifications. In brief: 5 ng of extracted DNA were used as template to amplify the V4 region of bacterial and archaeal 16S rRNA genes using universal primers 515F (5´-GTGYCAGCMGCCGCGGTAA-3´) and 806R (5´ GGACTACNVGGGTWTCTAAT-3´) (modified from Caporaso et al., 2011) and the platinum hot start PCR master mix (Invitrogen). Polymerase chain reaction (PCR) was conducted under the conditions of an initial denaturation of 94°C for 3 min followed by 10 cycles of denaturation at 94°C for 45 sec, annealing at 50°C for 60 sec, and elongation at 72°C for 90 sec. Subsequently, PCR products were purified using Agencourt AMPure XP magnetic beads (Beckman Coulter, Indianapolis, IN, United states), eluted in 24 µl PCR-grade water and 20 µl used as template for a second PCR with 14 cycles for the introduction of barcodes and Illumina sequencing adaptors. PCR conditions were the same as in the first PCR. After pooling equimolar amounts of template DNA, amplicon sequencing was performed at the Leibniz Institute DSMZ – German Collection of Microorganisms and Cell Cultures, Germany on the Illumina MiSeq platform using paired-end sequencing (2×300 bp, MiSeq reagents kit v3). Amplicon reads were processed using Qiime2 version 2023.5 [6], including quality control, amplicon sequencing variants (ASV) clustering and de novo chimera filtering using DADA2 [7]. Processed samples had on average 114,468 reads, with 90% of all samples having between 57,669 and 166,633 paired-end reads (5 and 95% quantiles, respectively). ASVs were taxonomically classified using the naïve Bayes classifier in Qiime2 and the Silva database v.138.1 [8].

### Supplementary Tables

**Supplementary Table 1.** Results of model selection for possible generalized additive models (gam) relating soil respiration to incubation time (day), bacterivorous nematode addition (treatment), and maize litter addition (substrate).

| Model | Modeled response under | Deviance expplained (%) | deltaAIC |
|---|---|---|---|
| mod0 <- gam(respiration ~ day) | no smooth. Null model. | 4.11 | 783.9 |
| mod_G <- gam(respiration ~ s(day)) | global smooth for day. | 32.2 | 672.0 |
| mod_GI <- gam(respiration ~ s(day) + s(day, by=treatment, bs=”tp”) | global smooth for day *plus* individual group smooths per day for nematode treatment. | 87.2 | 72.6 |
| mod_I <- gam(respiration ~ s(day, by=treatment, bs=”tp”) | individual group smooths per day for nematode treatment. | 87.2 | 72.6 |
| mod_GS <- gam(respiration ~ s(day) + s(day, by=treatment, bs=”fs”) | global smooth for day *plus* shared group smooths per day for nematode treatment. | 87.2 | 72.6 |
| mod_S <- gam(respiration ~ s(day, by=treatment, bs=”tp”) | shared group smooths per day for nematode treatment. | 87.2 | 72.6 |
| modS_ST <- gam(respiration ~ substrate + treatment + s(day, by=treatment, bs=”fs”) | additive effects of substrate addition, treatment type, and shared group smooths per day for nematode treatment. | 89.5 | 1.9 |
| modS_SintactT <- gam(respiration ~ substrate*treatment + s(day, by=treatment, bs=”fs”) | interacting effects of substrate addition and treatment type plus shared group smooths per day for nematode treatment. | 89.6 | 0.0 |

Diagnostic plots of the best fitting GAM (modS_SintactT) are given in Supplementary Figure 1.

**Supplementary Table 2.** Summary of the best-fitting GAM obtained used to explain soil respiration. The structure of the final model according to Supplementary Table 1 was modS_SintactT <- gam(respiration ~ substrate*treatment + s(day, by=treatment, bs="tp"); method = REML, family = Gaussian.

*Parametric coefficients*

| | Estimate | Std. error | t-value | *p*-value | Significance |
|---|---|---|---|---|---|
| Intercept | 1.36421 | 0.03331 | 40.951 | $< 2 \times 10^{-16}$ | *** |
| Substrate addition | 0.11751 | 0.02338 | 5.025 | $8.05 \times 10^{-07}$ | *** |
| Nematode treatment | 0.06989 | 0.04362 | 1.602 | 0.1100 | |
| Substrate:Nematode | 0.06661 | 0.03413 | 1.952 | 0.0518 | . |

Significance codes: 0 '***' ; 0.001 '**' ; 0.01 '*' ; 0.05 '.' ; 0.1 ' ' ; 1

*Approximate significance of smooth terms*

| | edf | Ref.df | F | *p*-value | Significance |
|---|---|---|---|---|---|
| s(day,treatment) | 15.74 | 16 | 170.2 | $< 2 \times 10^{-16}$ | *** |

Significance codes: 0 '***' ; 0.001 '**' ; 0.01 '*' ; 0.05 '.' ; 0.1 ' ' ; 1

$R^2$ (adj) = 0.89, deviance explained = 89.6%

-REML = -90.74, scale est. = 0.026648, n = 369

**Supplementary Table 3.** Analysis of variance (ANOVA) of total abundance, alpha diversity ($^0D$ and $^2D$), and rank abundance structure (alpha-gambin value) changes over time. N – bacterivorous nematode; M – maize litter. Significant results at $p < 0.05$ are given in bold.

| Parameter | Treatment | $F_{4,10}$-value | *p*-value |
|---|---|---|---|
| Total bacterial and archaeal 16S rRNA genes/g dry soil | **+N/+M** | **4.780** | **0.048** |
| | +N/–M | 0.316 | 0.583 |
| | –N/+M | 2.029 | 0.178 |
| | –N/–M | 0.001 | 0.907 |
| ASV richness ($^0D$) | +N/+M | 0.815 | 0.383 |
| | +N/–M | 1.370 | 0.262 |
| | –N/+M | 0.662 | 0.431 |
| | –N/–M | 0.632 | 0.441 |
| Dominant ASVs ($^2D$) | +N/+M | 0.403 | 0.537 |
| | +N/–M | 1.290 | 0.277 |
| | –N/+M | 0.127 | 0.727 |
| | –N/–M | 1.530 | 0.238 |
| Alpha-gambin | +N/+M | 0.042 | 0.840 |
| | **+N/–M** | **8.450** | **0.012** |
| | –N/+M | 0.015 | 0.903 |
| | **–N/–M** | **6.370** | **0.025** |

**Supplementary Table 4.** Post-hoc comparisons of group means for significant ANOVA results shown in Supplementary Table 3. Post-hoc tests were done using the R package multcomp.

*Total bacterial and archaeal 16S rRNA genes (g dry soil)$^{-1}$, Treatment +N/+M*

| Linear Hypotheses | Estimate | Std. Error | t value | Adjusted p-value |
|---|---|---|---|---|
| 4 - 0 == 0 | 3.61E+07 | 2.27E+07 | 1.590 | 0.501 |
| 8 - 0 == 0 | 5.88E+07 | 5.18E+07 | 1.136 | 0.760 |
| 16 - 0 == 0 | -5.42E+07 | 1.95E+07 | -2.780 | 0.096 |
| 32 - 0 == 0 | -3.82E+07 | 2.15E+07 | -1.773 | 0.405 |
| 8 - 4 == 0 | 2.27E+07 | 5.00E+07 | 0.453 | 0.988 |
| **16 - 4 == 0** | **-9.03E+07** | **1.42E+07** | **-6.351** | **0.001** |
| **32 - 4 == 0** | **-7.43E+07** | **1.69E+07** | **-4.391** | **0.008** |
| 16 - 8 == 0 | -1.13E+08 | 4.86E+07 | -2.323 | 0.192 |
| 32 - 8 == 0 | -9.70E+07 | 4.95E+07 | -1.960 | 0.319 |
| 32 - 16 == 0 | 1.60E+07 | 1.23E+07 | 1.305 | 0.665 |

*Alpha-gambin, Treatment +N/-M*

| Linear Hypotheses | Estimate | Std. Error | t value | Adjusted p-value |
|---|---|---|---|---|
| 4 - 0 == 0 | 2.88E-03 | 0.200 | 0.014 | 1 |
| 8 - 0 == 0 | -1.43E-01 | 0.184 | -0.775 | 0.924 |
| 16 - 0 == 0 | -1.76E-01 | 0.182 | -0.965 | 0.851 |
| 32 - 0 == 0 | -2.67E-01 | 0.175 | -1.530 | 0.542 |
| 8 - 4 == 0 | -1.46E-01 | 0.122 | -1.199 | 0.732 |
| 16 - 4 == 0 | -1.79E-01 | 0.119 | -1.509 | 0.554 |
| 32 - 4 == 0 | -2.70E-01 | 0.106 | -2.541 | 0.143 |
| 16 - 8 == 0 | -3.32E-02 | 0.090 | -0.369 | 0.995 |
| 32 - 8 == 0 | -1.25E-01 | 0.073 | -1.702 | 0.448 |
| 32 - 16 == 0 | -9.14E-02 | 0.068 | -1.341 | 0.651 |

*Alpha-gambin, Treatment -N/-M*

| Linear Hypotheses | Estimate | Std. Error | t value | Adjusted p-value |
|---|---|---|---|---|
| 4 - 0 == 0 | -2.07E-01 | 0.196 | -1.058 | 0.811 |
| 8 - 0 == 0 | -4.26E-01 | 0.248 | -1.718 | 0.446 |
| 16 - 0 == 0 | -2.60E-01 | 0.334 | -0.780 | 0.925 |
| 32 - 0 == 0 | -5.75E-01 | 0.208 | -2.770 | 0.104 |
| 8 - 4 == 0 | -2.19E-01 | 0.187 | -1.172 | 0.751 |
| 16 - 4 == 0 | -5.30E-02 | 0.291 | -0.182 | 1 |
| 32 - 4 == 0 | -3.68E-01 | 0.128 | -2.869 | 0.089 |
| 16 - 8 == 0 | 1.66E-01 | 0.328 | 0.505 | 0.983 |
| 32 - 8 == 0 | -1.49E-01 | 0.199 | -0.749 | 0.934 |
| 32 - 16 == 0 | -3.15E-01 | 0.299 | -1.053 | 0.813 |

**Supplementary Table 5.** Overview over abundance changes, taxonomic affiliation, and response type of dominant ASVs under the different treatment combinations. M – maize litter , N – nematode, + and – signs indicate presence and absence, respectively. The file is provided by a separate Excel file.

#### Supplementary Figures

**Supplementary Figure 1.** Overview of experimental design. The graph was created with BioRender.com under license number LI26OANQ3V for use in journal publications.

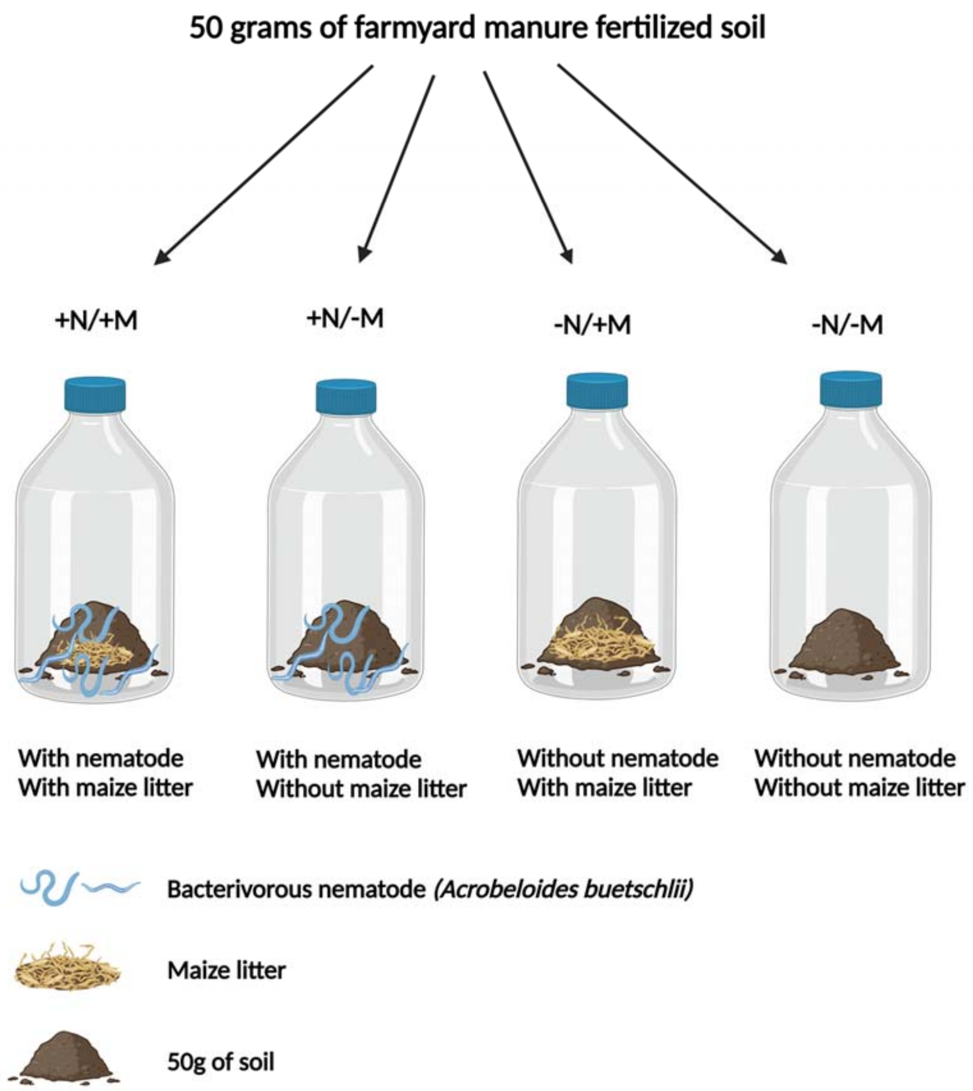

**Supplementary Figure 2.** Diagnostic plots of the best fitting GAM (modS_SintactT) used to explain soil respiration in relation to day, treatment, and substrate

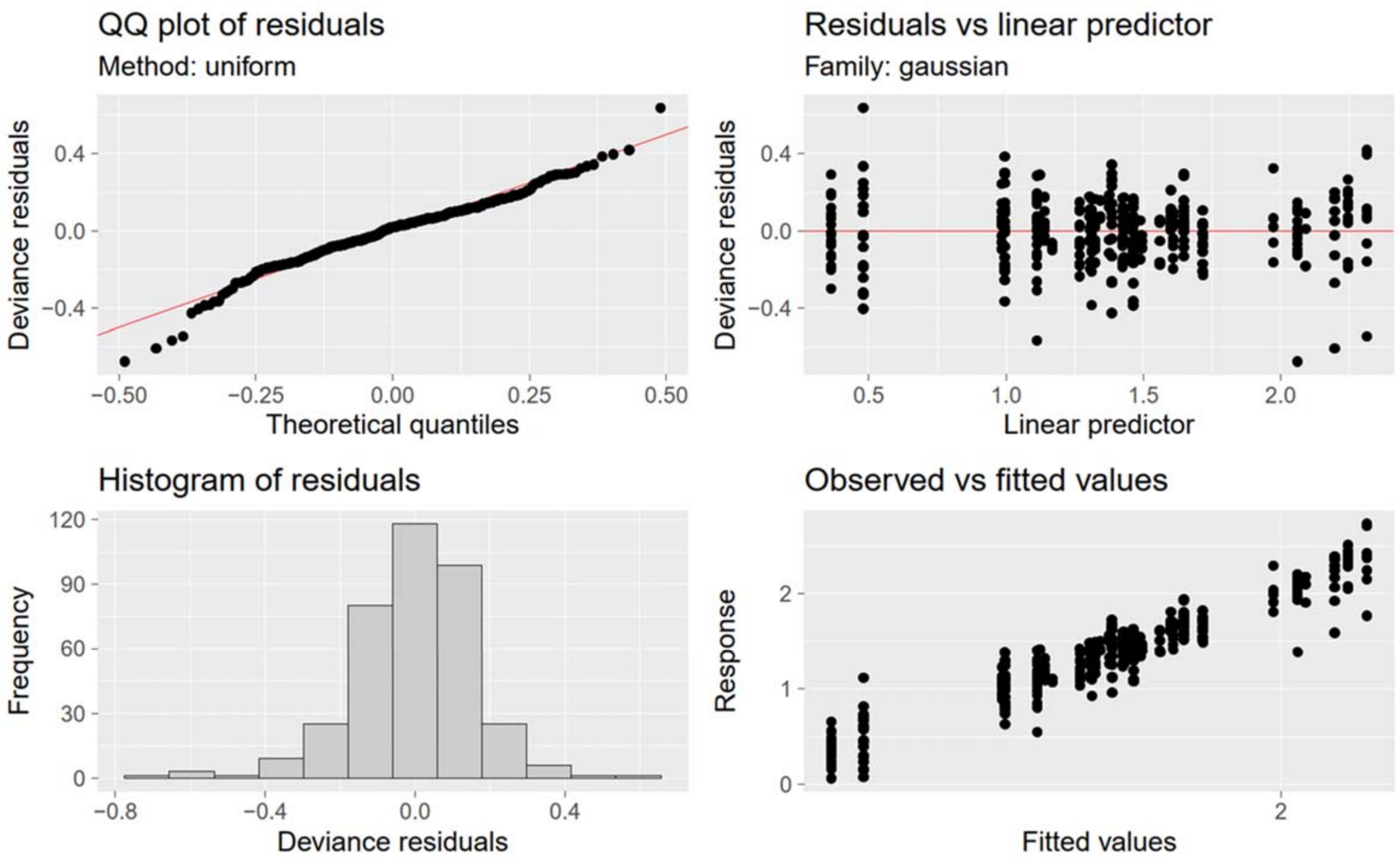

**Supplementary Figure 3.** Absolute abundance and alpha diversity of total Bacteria and Archaea throughout the experiment. Alpha diversity was based on Hill number analysis [9, 10]. The alpha-gambin value is provided as a parameter describing rank abundance distributions [11].

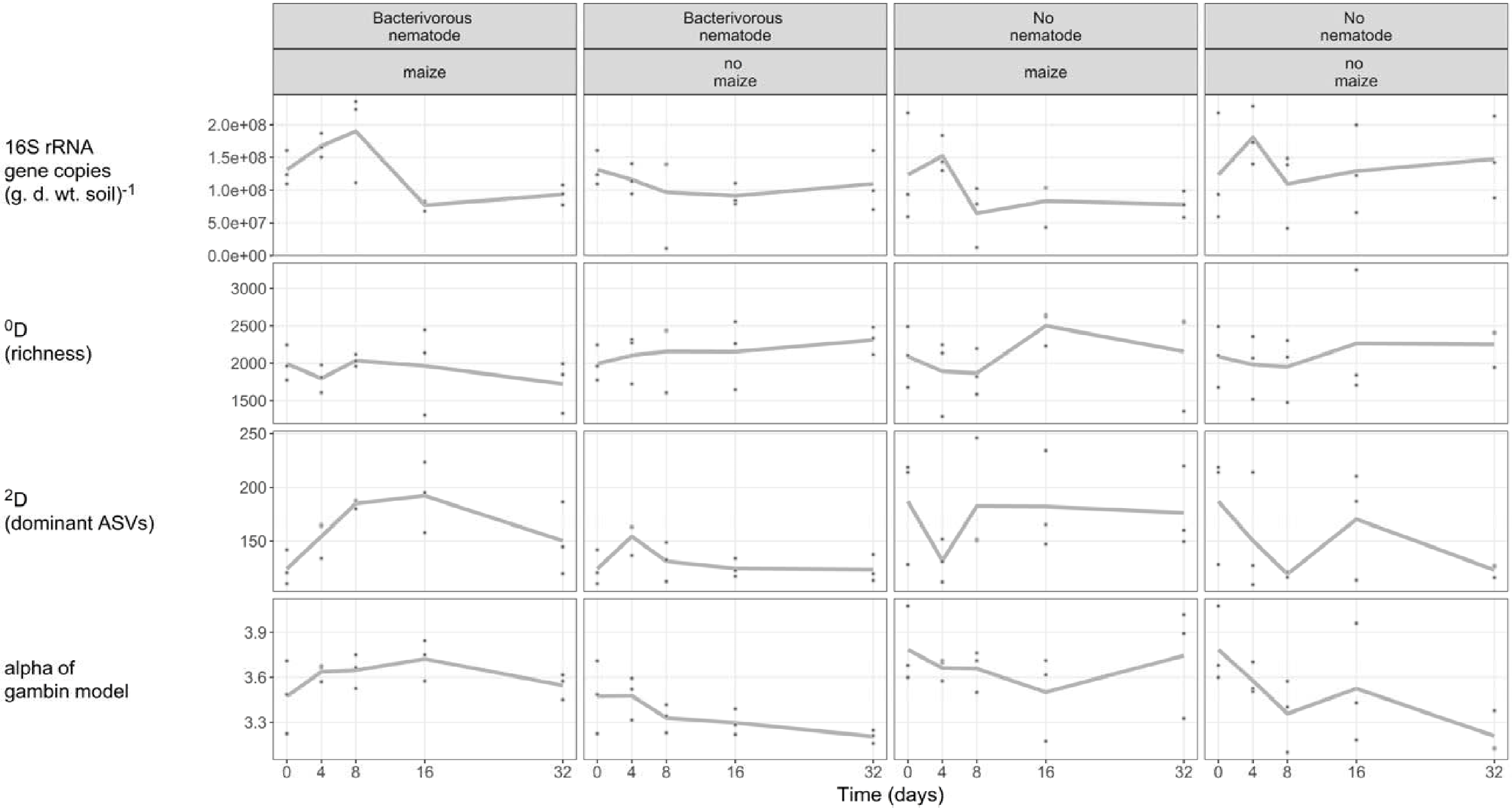

**Supplementary Figure 4.** Time-resolved beta diversity changes of the bacterial and archaeal community in response to addition of the bacterivorous nematode *A. buetschlii* (indicated in the panels) and maize litter (+M – added maize litter; −M – no addition of maize litter). The analysis was based on non-metric multidimensional scaling (NMDS) of weighted unifrac (left side) and Bray-Curtis (right side) distance metrics.

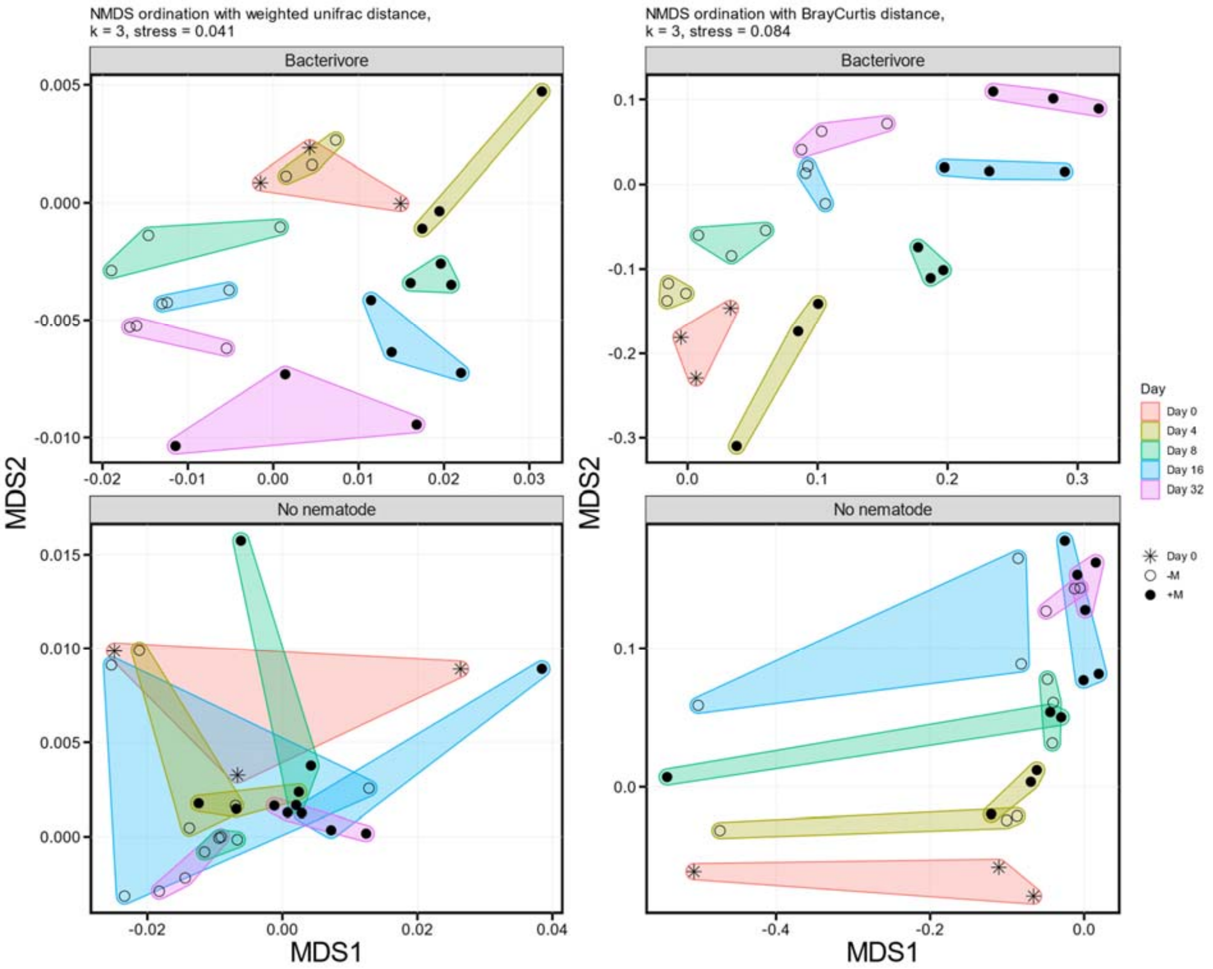

**Supplementary Figure 5.** Changes of beta diversity metrics over time in response to addition of the bacterivorous nematode *A. buetschlii* and maize litter. Beta-diversity distances were calculated between all pairs of replicates. As different distance estimates (weighted unifrac versus Bray-Curtis) may yield distance values at different scales, all distance values from a specific distance measure were scaled to mean = 0 and standard deviation = 1. Each boxplot represents nine pairwise comparisons (three replicates of one time point compared to all three replicates of another time point). Only the boxplot at time point 0 is covered by 15 data points (three pairs of replicate comparison per time point, five time points).

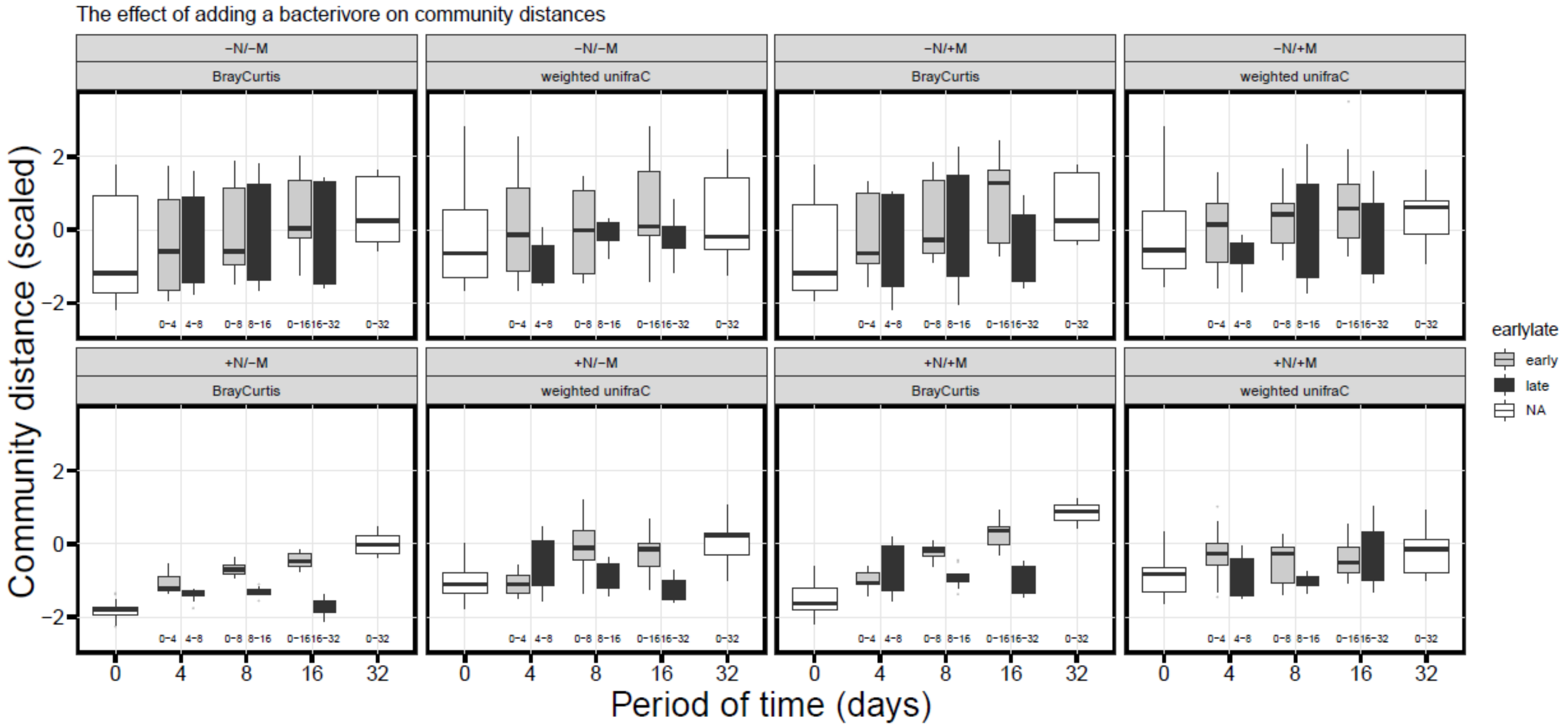

**Supplementary Figure 6.** Rank abundance distribution of all observed response type patterns as based on the summed median abundances of their representing ASVs. Response types of individual ASVs are shown in Supplementary Table 5. Letters correspond to response types shown in Fig. 3. If representing ASVs did not belong to a certain response type under a given treatment "_" is indicated. The sequence of letters corresponds to treatments in the following order: +N/+M, +N/−M, −N/+M, −N/−M.

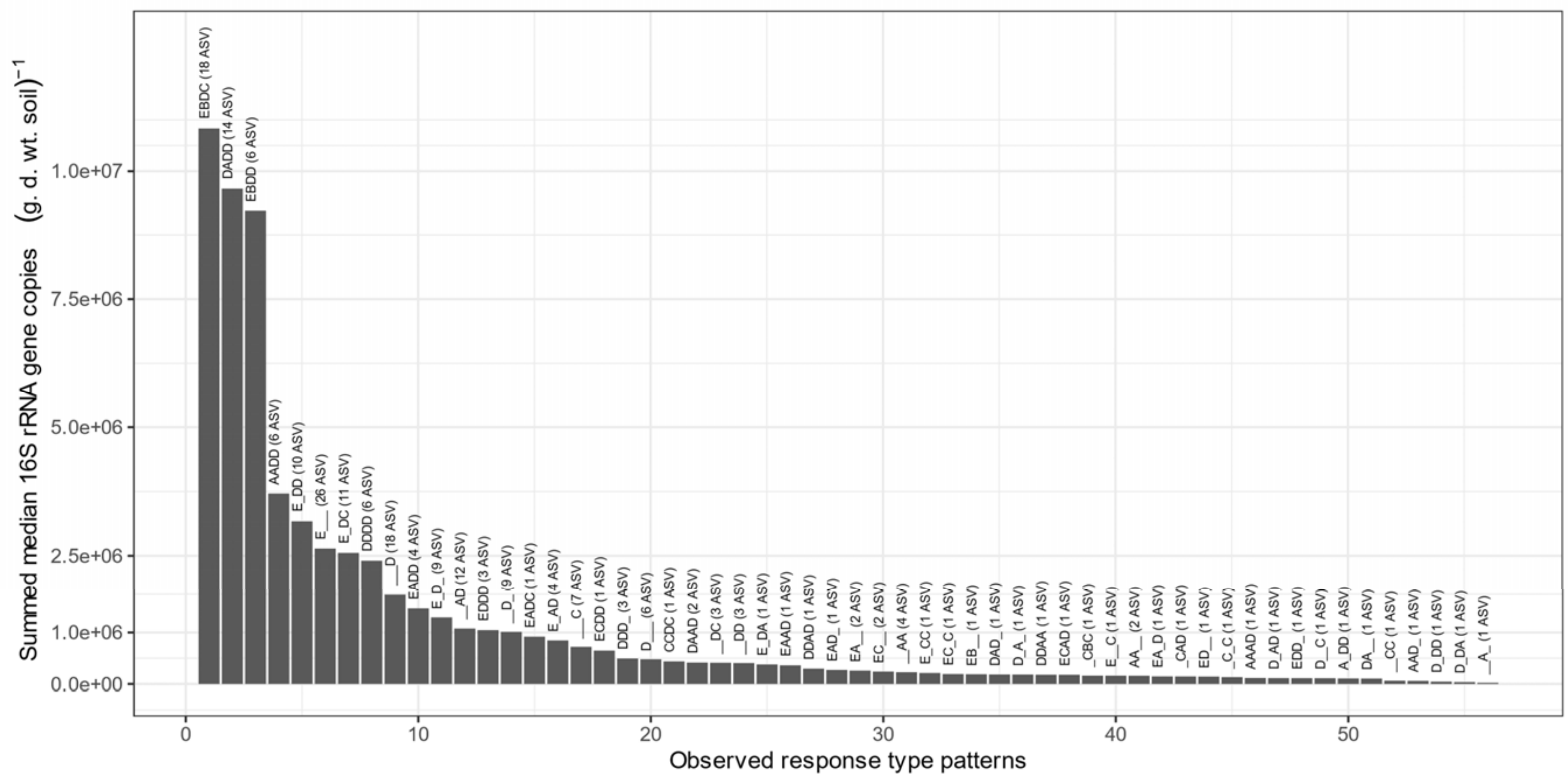
